# Amplification of the epigenetic (gestational) age acceleration signal

**DOI:** 10.1101/2021.02.02.429312

**Authors:** Yunsung Lee, Jon Bohlin

## Abstract

**Background:** Epigenetic (gestational) age acceleration (E(G)AA) is associated with environmental exposures and health outcomes in humans. However, E(G)AA is the residual term from a regression of epigenetic age (outcome) on chronological (gestational) age (predictor) and therefore strongly obscured by ‘noise’ from multiple sources. Here, we propose a simple procedure, based on regression, principal component analysis (PCA), and the Lasso, that amplifies E(G)AA signals. More specifically, we first regress given (gestational) age against each CpG used for epigenetic (gestational) age prediction. The CpGs are typically taken from one of several epigenetic clocks available. PCA is subsequently performed on the resulting matrix of residual vectors for each CpG as it projects the E(G)AA signal onto perpendicular principal components (PCs), thereby separating ‘signal’ from noise. Finally, we use the Lasso to select PCs associated with an outcome of interest. We apply our method to previous studies: EAA in patients with Down’s syndrome and Werner’s syndrome and EGAA of newborns exposed to prenatal smoking as well as associations with maternal BMI.

**Results:** The extracted EAA components computed using our proposed procedure revealed a significant association with Down’s syndrome (P_B_<0.05, Bonferroni adjusted for multiple testing) as well as for Werner’s Syndrome (P_B_<0.05). For EGAA we find a significant association with maternal prenatal smoking (P_B_<0.05, also Bonferroni adjusted) and maternal BMI (P_B_<0.05). Additionally, by examining the loadings of the PCs of interest, and contrary to residual EGAA, our method can identify implicated CpGs.

**Conclusions:** Our findings suggest that our proposed procedure leads to a remarkable amplification of the E(G)AA signal. Furthermore, our method reveals that E(G)AA is a composite signal that can be driven by multiple independent factors.

## BACKGROUND

Epigenetic age (EA) has been found to reflect biological aging in humans [1]. Ever since epigenetic age estimators (often referred to as epigenetic clocks) were first introduced [2–4] many attempts have been made to link EA to longevity and various diseases or exposures. For example, higher EA compared with chronological age (CA) was associated with all-cause mortality in later life [5], HIV [5, 6], early menopause [7], Parkinson’s disease [8], Hutchinson Gilford Progeria Syndrome [9], Alzheimer’s disease [10], Huntington’s disease [11], brain-degenerative disorders [12] and more [13, 14]. Furthermore, patients with Down’s syndrome [15] and Werner’s syndrome [16] also exhibit high EA with respect to CA. Lower EA with respect to CA has, on the other hand, been found in centenarians and been associated with longevity [17].

Analogously, epigenetic gestational age (EGA) differing from ultrasound/last menstruation period-based estimates of gestational age (GA) in newborns has also been associated with a set of adverse conditions [18, 19].

Both increased and decreased EA compared with CA are usually referred to as epigenetic age acceleration (EAA). It is defined as the residual term from a regression of EA on CA. However, by its construction EAA, as well as the analogous epigenetic gestational age acceleration (EGAA), is considerably influenced by ‘noise’ generated from multiple sources such as CpG probes, measurement error, dataset normalization procedures and the regression model itself [20]. We here suggest a simple procedure to extract and amplify EAA (or EGAA) signals from ‘noise’. We demonstrate our procedure on 1) EAA with respect to Down’s syndrome and Werner’s syndrome [15] 2) EGAA with respect to prenatal maternal smoking and association with maternal BMI [18, 21]. We also compare how our method performs with respect to standard residual E(G)AA.

## RESULTS

### The procedure

To boost the E(G)AA signal, we first extract the CpGs associated with epigenetic (gestational) age prediction. A linear regression is performed for each CpG with CA (or GA) as the outcome. The residuals from each regression model are placed in a matrix on which a subsequent principal component analysis (PCA) is performed. Finally, a Lasso regression [22] is carried out with the phenotype of interest as the outcome, e.g., Down’s syndrome and maternal BMI, and the principal components (PC) as the feature matrix. The effects of selected PCs and phenotype of interest can subsequently be assessed using standard regression analysis. The CpGs associated with E(G)AA may be interrogated further by examining the corresponding PCA loadings. We provide the specific details of the procedure in the Methods section as well as in a separate R script included as Additional file 1.

### Revisited literature and study samples

We first revisited the study “Accelerated epigenetic aging in Down syndrome” [15]. This study was selected for the two following reasons: 1) The association between EAA and Down syndrome is already considered to be solid, and 2) the DNAm data used in the article is publically available (GSE52588, Table 1). We considered four epigenetic clocks: Hannum et al.’s clock [2], Horvath’s Pan-Tissue clock [3], Levine et al.’s PhenoAge clock [23] and Horvath skin and blood clock [9], all of which can be applicable to GSE52588 (Figures 1 and 2). In addition, we also re-examined a study on Werner’s syndrome [16] (GSE131752, Table 1) where we also consider the same epigenetic clocks as mentioned above for Down’s syndrome (Additional file 2, Figures S1 and S2).

**Table 1.**
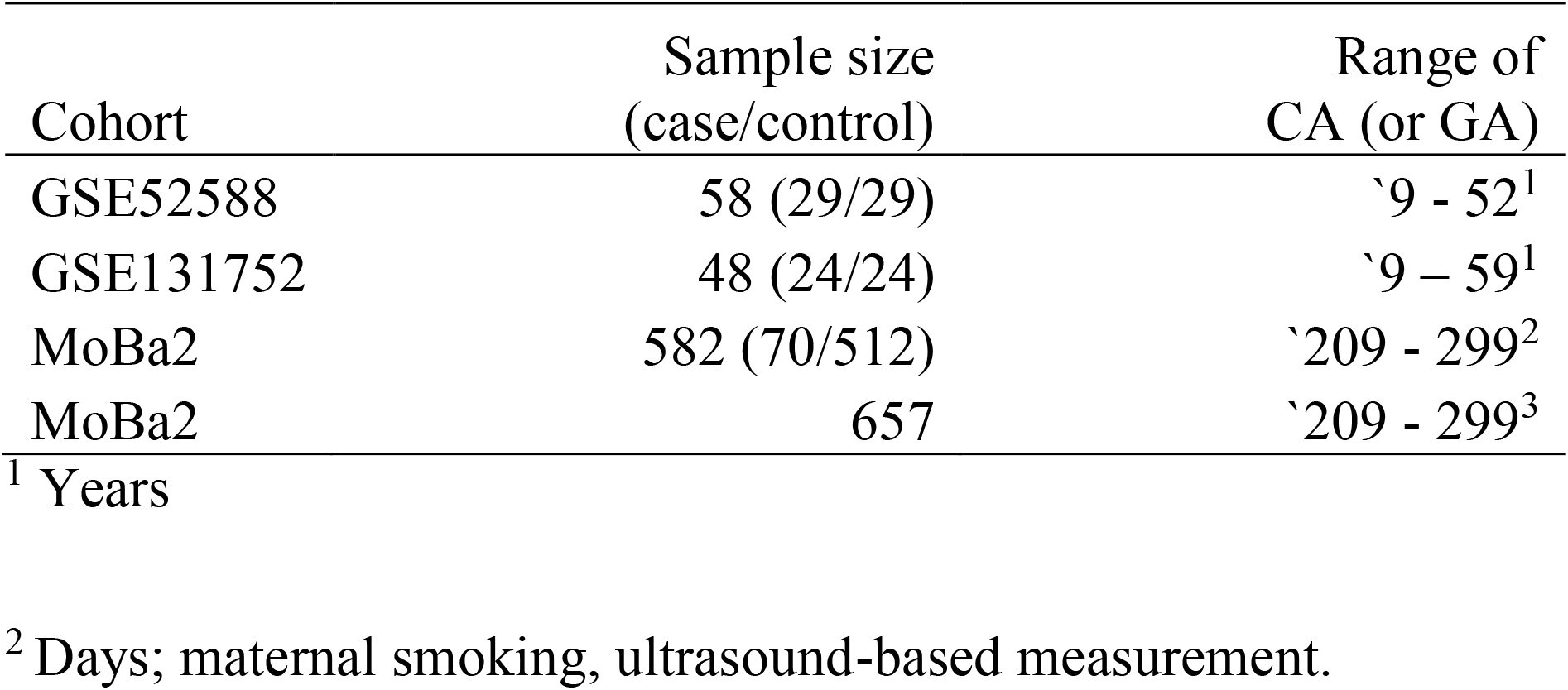
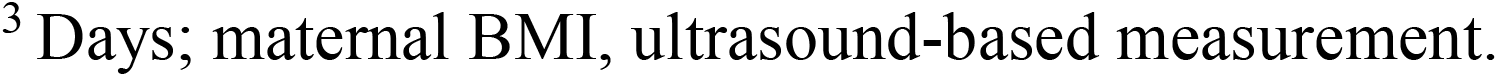
Descriptive statistics of study samples.

| Cohort | Sample size<br>(case/control) | Range of<br>CA (or GA) |
| --- | --- | --- |
| GSE52588 | 58 (29/29) | `9 - 52<sup>1</sup> |
| GSE131752 | 48 (24/24) | `9 – 59<sup>1</sup> |
| MoBa2 | 582 (70/512) | `209 - 299<sup>2</sup> |
| MoBa2 | 657 | `209 - 299<sup>3</sup> |

<sup>1</sup> Years
<sup>2</sup> Days; maternal smoking, ultrasound-based measurement.
<sup>3</sup> Days; maternal BMI, ultrasound-based measurement.

**Figure 1.**
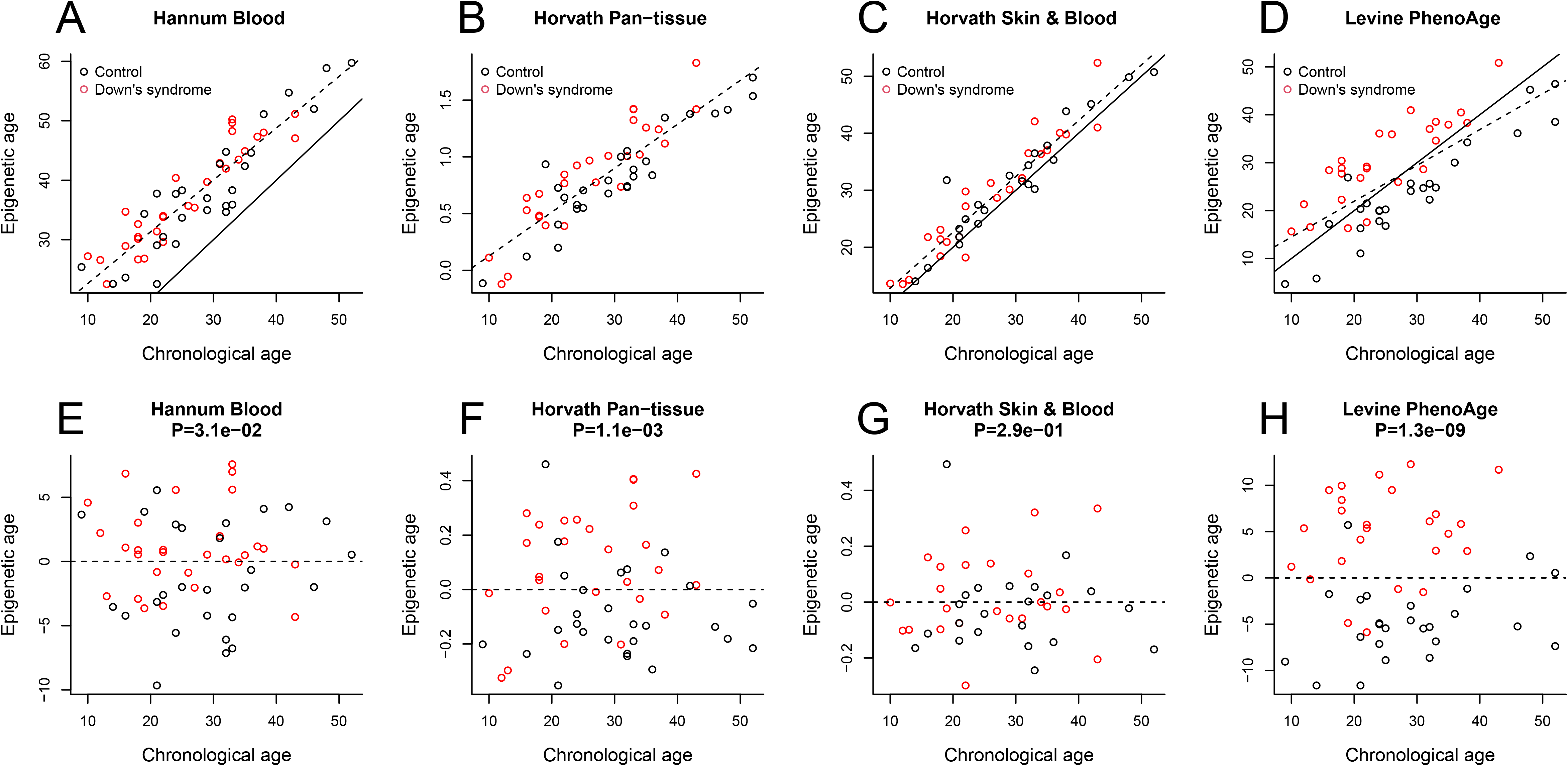
Associations between residual EAA and Down syndrome. **A-D:** Scatter plot of CA versus EA. **E-H:** Scatter plot of CA versus EAA. The black dots refer to unaffected controls whereas the red dots refer to subjects with Down syndrome. P-values were obtained from student t tests comparing EAAs between controls and subjects with Down syndrome.

**Figure 2.**
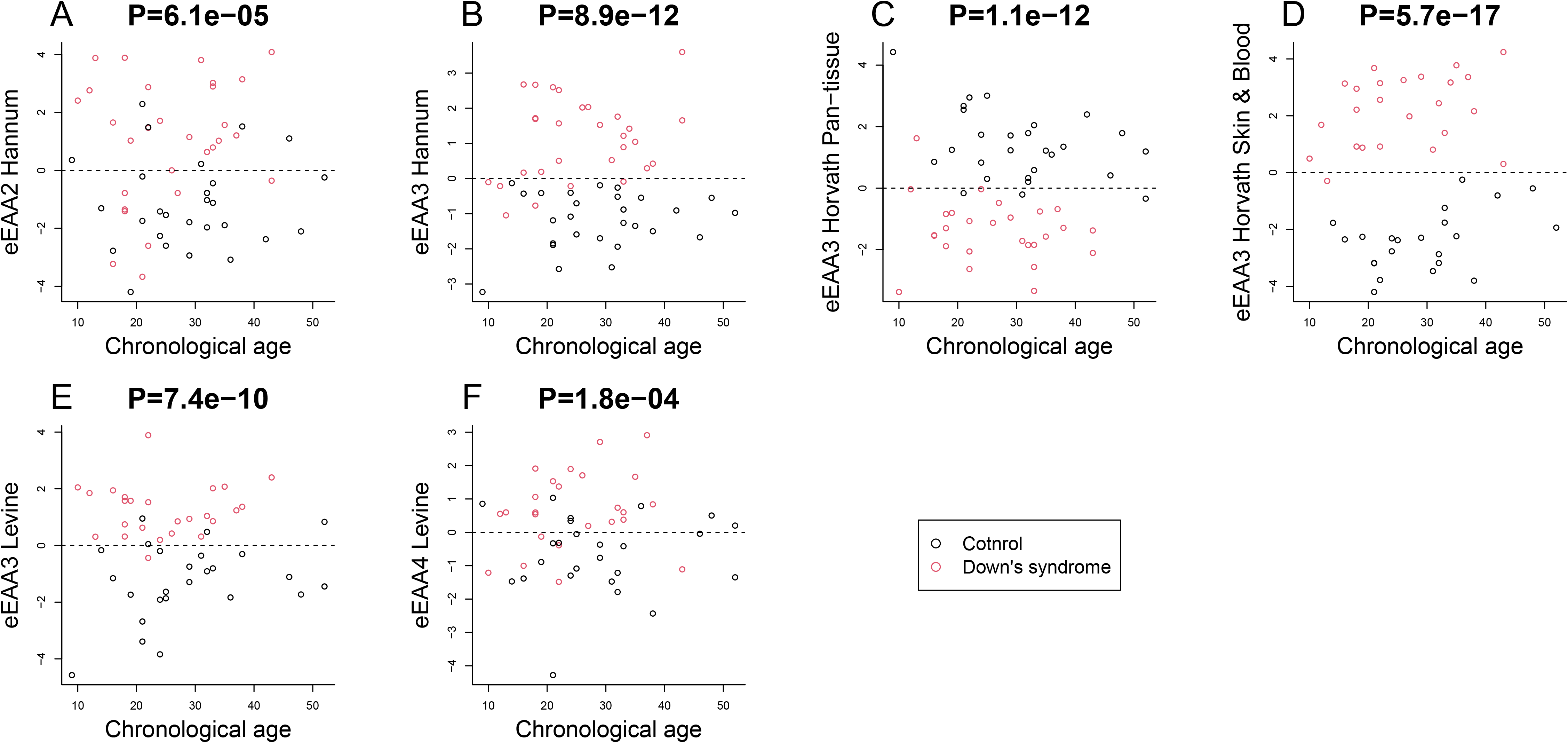
Associations between eEAAs and Down’s syndrome. **A:** eEAA2 (i.e., PC2) obtained by Hannum et al.’s clock. **B:** eEAA3 (i.e., PC3) obtained by Hannum et al.’s clock. **C:** eEAA3 (i.e., PC3) obtained by Horvath’s Pan-Tissue clock. **D:** eEAA3 (i.e., PC3) obtained by Horvath et al.’s Skin & Blood clock. **E:** eEAA3 (i.e., PC3) obtained by Levine et al.’s PhenoAge clock. **F:** eEAA4 (i.e., PC4) obtained by Levine et al.’s PhenoAge clock. The black dots refer to unaffected controls whereas the red dots refer to subjects with Down syndrome. P-values were obtained from student t tests comparing eEAAs between controls and subjects with Down syndrome.

Secondly, we tested our EGAA-extraction method on cord blood DNAm, taken from newborns from the MoBa cohort study (MoBa2, n=685, Table 1), to assess EGAA previously reported with regards to maternal BMI and maternal smoking [18, 21].

### Case 1: Extracted EAAs in adults with Down’s- and Werner’s syndrome

We tested whether residual EAA differs between unaffected controls (*n*=29) and patients with Down syndrome (*n*=29). Although the original study [15] used Horvath’s Pan-Tissue clock only, we additionally tested other epigenetic clocks, e.g., Levine et al.’s PhenoAge clock and Horvath et al.’s Skin & Blood clock. The residual EAA from all clocks except Horvath’s Skin & Blood clock was significantly higher in subjects with Down syndrome than in unaffected controls (Figure 1).

Our PCA based EAA extraction method was applied to the same dataset (Details of the EAA extraction method can be found in the Methods section). The Lasso method extracted several PCs (we refer to these as eEAAs), and we tested the differences in the PCs between patients with Down’s syndrome and controls. In Figure 2, we demonstrate this using two eEAAs (eEAA2 and eEAA3, i.e., the second and third PC) from Hannum et al.’s Blood clock, one eEAA (eEAA3) from Horvath’s Pan-Tissue clock, eEAAs (eEEA3) from Horvath’s Skin & Blood clock, and two eEAAs (eEEA3 and eEEA4) from Levine’s PhenoAge clock, all of whom were found to be significantly associated with Down’s syndrome (P_B_<0.05) after the Bonferroni correction. Strikingly, the associations between eEAAs and Down’s syndrome resulted in considerably lower p-values than the associations between EAA and Down’s syndrome (compare Figure 2 with Figure 1). In addition, the fact that several independent eEAAs were found to be significantly associated with Down’s syndrome suggests that EAA is a multi-factorial composite trait. Our method also showed similar results for data on Werner’s syndrome [16, 24] (GSE131752), where the associations of Werner’s syndrome with standard EAAs were found to be substantially weaker than those with eEAAs resulting from our PC-based method (P_B_<0.05, Figures S1 and S2, Additional file 2).

### Case 2: Extracted EGAAs in newborns exposed to prenatal smoking and maternal BMI

In MoBa2 (n=685), we associated residual EGAA with maternal smoking during pregnancy. The standard residual-based EGAA was defined as the residuals from a regression of EGA on GA that was estimated by Bohlin et al.’s epigenetic GA clock [25]. We could not detect any significant association between residual EGAA and prenatal smoking in MoBa2 (P=0.18), while Girchenko et al. did [18] (see however a comment on this study by Simpkin et al. [26]).

We then applied our E(G)AA-extraction method and associated the resulting eEGAAs with prenatal smoking (Figure 3). We found that eEGAA9 and eEGAA90 were significantly associated (P_B_<0.05, Bonferroni corrected) with prenatal smoking. Since effects from prenatal maternal smoking on newborns’ methylome have been studied extensively we interrogated the loadings for eEGAA9 and eEGAA90 to see whether any CpG’s overlapped with those found in a meta-analysis [27]. In total we found 15 overlapping CpG’s of varying effect sizes (See Additional file 3).

**Figure 3.**
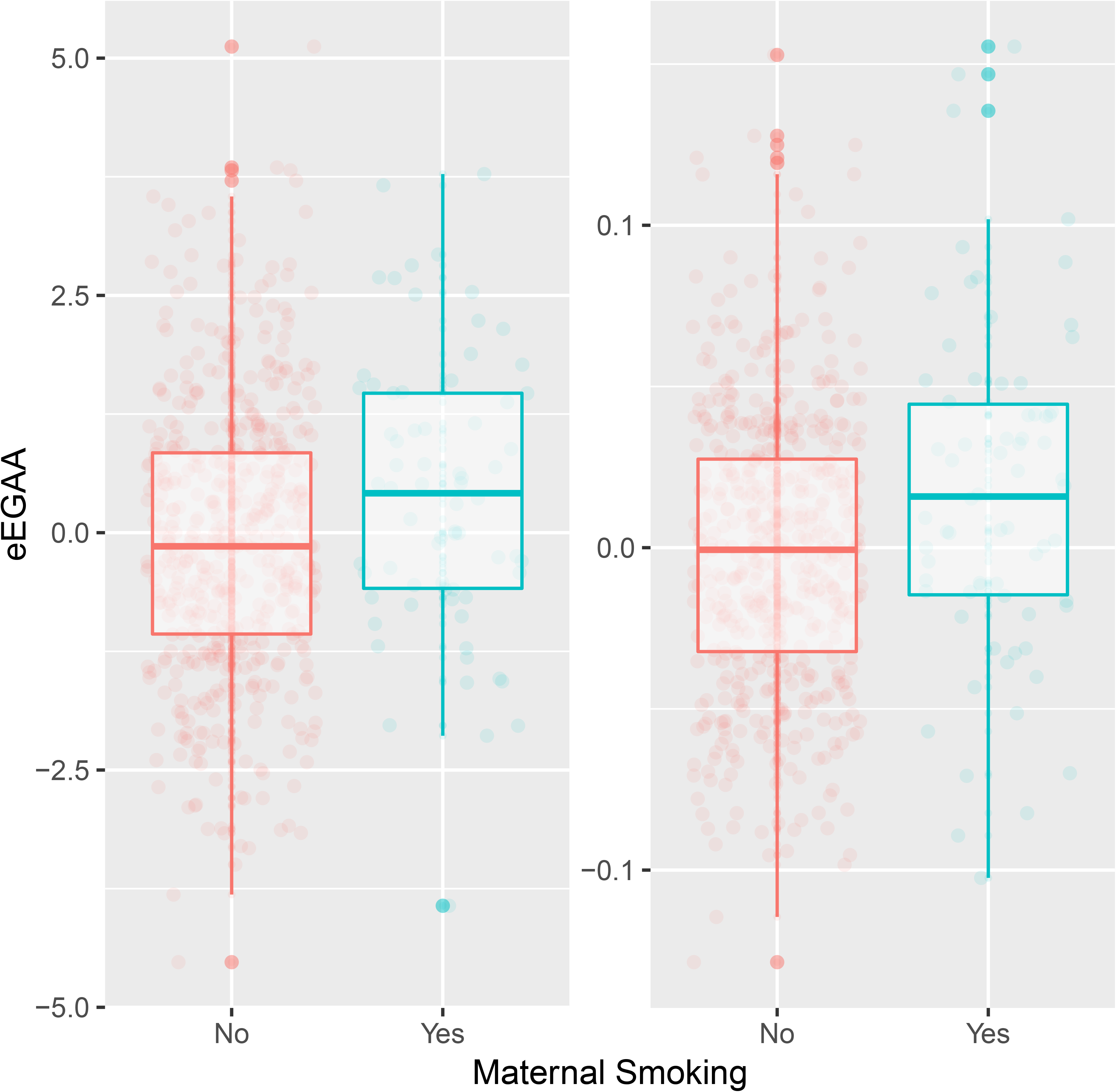
Association between eEGAAs and prenatal smoking. Box plots to contrast eEGAA9 (left) and eEGAA90 between unexposed newborns (i.e., non-smoker) and newborns exposed to prenatal smoking (i.e., smoker).

We also found a statistical association between eEGAA5 and maternal BMI (P_B_<0.05, Figure 4) but not for residual EGAA (P=0.85). Two studies have previously reported the association between maternal BMI and EGAA [18, 21].

**Figure 4.**
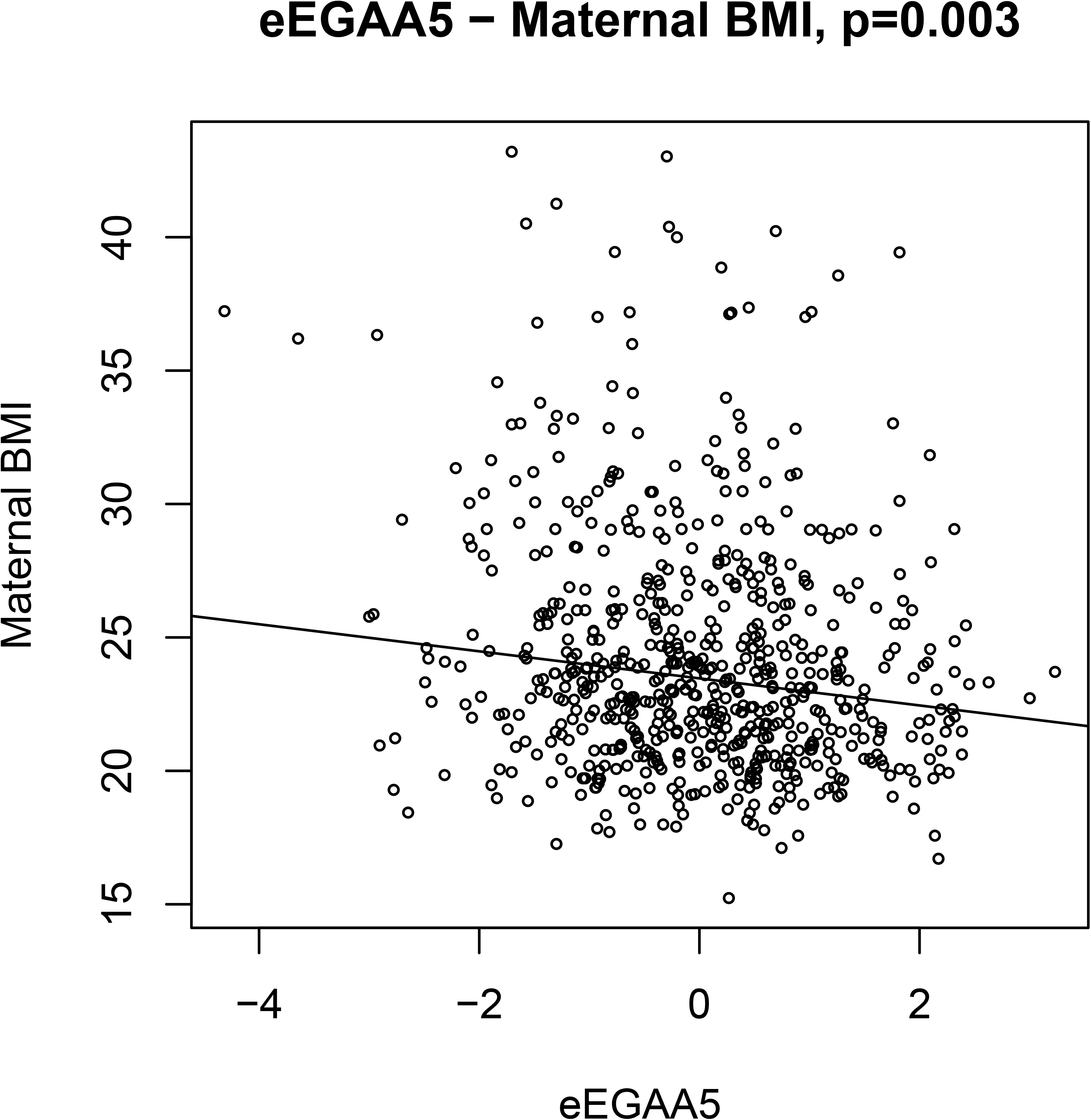
Association between eEGAA5 and maternal BMI. The figure shows maternal BMI (vertical axis) plotted against eEGAA5 (horizontal axis).

## DISCUSSION

Our new procedure amplifies eE(G)AA signals as compared with previous findings on residual E(G)AA. The core of this method lies in its micro-level approach which focuses on associating CA with each of the CpGs included in epigenetic clocks, separately (Additional file 4, Figure S4). The method thereby breaks the composite residual E(G)AA signal into multiple different independent components using PCA. Expectedly, in many of the CpG-CA relationships, subjects with a medical condition (e.g., Down’s syndrome, Figures 1–2, Werner’s syndrome, Additional file 2, Figure S1-S2) formed a shifted cluster [28]. This shift suggests that the subjects with Down’s and Werner’s syndrome biologically aged faster than the unaffected controls. When this shift was present, our regression of CA on a CpG amplified the difference between controls and subjects with Down’s and Werner’s syndrome by assigning mostly positive residuals to controls and negative residuals to Down’s- and Werner’s syndrome subjects, respectively (Figure 2 and Figure S2 in Additional file 2). However, this shift was not necessarily observed for all CpG-CA relationships; our procedure exclusively selected CpG-CA relationships where the shift occurred and generated PCs accordingly. As PCA projects variance explained from the covariates onto perpendicular components that are rotated across the CpG feature space, it may enable identifying CpGs responsible for E(G)AA, which are not easily discovered through a standard EWAS.

The conventional way of estimating residual EAA includes, as mentioned above, ‘noise’ from various sources. First, DNAm measurement at each CpG contains a substantial fraction of noise due to the way the signal is measured [29]. Second, DNAm array normalization methods may introduce additional bias and noise [30]. Third, the regression of EA on chronological age also introduce an additional noise term. Moreover, the link between residual E(G)AA and a phenotype of interest may be obscured by the quality of the clock CpGs used (see Figure 1); on the other hand, epigenetic clocks that show highly accurate CA predictions might not give any room for E(G)A to differ among the groups as was seen with Horvath’s Skin & Blood clock in Figure 1. Our E(G)AA-extraction method addresses these sources of variation by employing PCA. This means that independent noise, errors and variation in the data are projected onto separate components, while EAA associated CpGs end up with the same component resulting in an amplified signal. As such, our method could potentially also facilitate discovery in standard EWAS analyses between phenotypes and numerous CpGs with small effects that would not have been identified separately.

One general weakness with the Lasso is that it can select covariates that are associated amongst themselves but less so with the outcome [31]. However, since we use the PCA matrix as the feature matrix (i.e. covariate matrix), this weakness is avoided as principal components are uncorrelated. Hence, our described method only extracts features (i.e. principal components) that are associated with the outcome of interest. Since the PCs (or eE(G)AAs) are uncorrelated, adjustment for multiple testing must be performed using the Bonferroni method.

Unlike residual E(G)AA, our method suggests that E(G)AA may exist in multiple forms. As presented in Figure 2, we found that several independent eEAAs (i.e., PCs) were associated with Down’s syndrome. The fact that eE(G)AAs are orthogonal to one another indicates that there can be several independent and uncorrelated mechanisms influencing E(G)AA. The standard residual-based E(G)AA measure [5] is intuitive due to its simple estimation of E(G)AA. Regrettably, the estimation of residual based E(G)AA restricts analyses to one bulk estimate of E(G)AA incorporating several sources of signals and noise. Our method, however, can identify the individual contributions of CpGs to each eE(G)AA. By the property of PCA, eE(G)AAs are linear combinations of residuals (from a linear regression of CA on each CpG) and loadings. Associations between CpGs and eE(G)AA can therefore be interrogated through exploration of genomic contexts such as genes, gene bodies, gene networks, genomic positions and genomic regions, e.g., CpG islands, shores, shelfs, and promoter regions.

Our proposed procedure also has some limitations; first, our method might need adjustment for batch effects or cell type compositions [32, 33]. A poor study design separating cases and controls may result in amplified E(G)AA signal differences between these groups. However, this is also a challenge for other measures and conventional EWAS analyses and therefore not specific to our described method. Second, it can be difficult to determine the sign (direction) of eE(G)AAs since the PCs are rotated independent of the outcome. The last limitation is the reason we primarily consider our method to be an E(G)AA signal boosting method, although it can readily identify novel associations not detectable with residual E(G)AA methods, as was shown for EGAA above.

Finally, we would like to emphasize that our procedure is not strictly dependent on epigenetic clocks. Although epigenetic clocks have been known for excellent performance in selecting chronological age (or GA) related CpGs that maximize prediction accuracy [14, 26], an EWAS can also select chronological age (or GA) related CpGs. Hence, our proposed method could potentially be of utility also for EWAS-based studies where noisy probes may hamper interpretation of results.

## CONCLUSIONS

We demonstrated a simple procedure, using linear regression, principal component analysis and the Lasso, that extracts E(G)AA from noise and identifies CpGs associated with E(G)AA. The method also indicates that the E(G)AA signals may consist of independent biological processes. While the mechanisms underlying E(G)AA are not known, this procedure should facilitate further exploration into epigenetic aging research.

## METHODS

### Description of the procedure

We assume CA (or GA) to be a linear combination of DNAm levels at *n* CpG sites as follows.

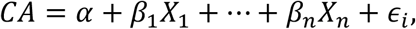

Where *X*_*j*_ is the DNAm level at the *j*-th CpG site, and *ϵ*_*i*_ is an error term assumed to be normally distributed with mean 0 and variance equal to *σ*^2^ (i.e.,*ϵ* ~ *N*(0, *σ*^2^)). We calculate EA (or EGA) using the estimated values of the *β*_1_,…, *β*_*n*_ provided by various epigenetic clocks.

We extract DNAm one CpG at a time and regress CA (or GA) on the extracted DNAm for each CpG using standard linear regression (we prefer the robust MM-type regressor [34]).

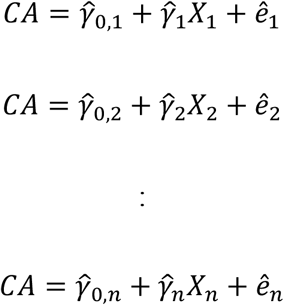

We assume that EA (or EGA) is a linear combination of DNAm levels at certain CpGs and this enables us to obtain *n* residuals (*ê*_1_,…, *ê*_*n*_) from the *n* different linear regressions. With these residual terms (*ê*_1_,…, *ê*_*n*_), we made a residual matrix ***R*** = [*ê*_1_, *ê*_2_,…, *ê*_*n*_] on which we applied PCA. This PCA resulted in a principal component (PC) matrix, ***P*** = [*PC*_1_, *PC*_2_,…, *PC*_*n*_], and a vector of variances, ***λ*** = [*λ*_1_, *λ*_2_,…, *λ*_*n*_]. We selected PCs associated with the outcome of interest by employing the Lasso [22] with the matrix ***P*** as the covariate/feature matrix and the phenotype of interest as the outcome y. Since the matrix ****P**** consists of perpendicular vectors further statistical inference should correct for multiple testing based on the number of residual terms in the matrix ***R***.

Here, we used Lasso (‘glmnet’ R package [35]) to select CpGs for E(G)AA. Further statistical inference can be performed using standard regression-based methods along with the Bonferroni adjustment for multiple testing. We emphasize that CpGs are not necessarily selected from epigenetic clocks but can be selected from a standard EWAS of CA (or GA).

All statistical analyses were carried out using the free software R v. 3.6.1 [36].

### Materials

GSE52588 is a publically available DNAm dataset consisting of 1) subjects with Down syndrome (*n*=29), 2) their unaffected siblings (*n*=29) and 3) unaffected mothers (*n*=29) [37]. As described in the results section, the subjects with Down syndrome and their siblings were selected for this study (*n*=58). DNAm was measured using the Illumina HumanMethylation450 BeadChip (Illumina, San Diego, CA, USA). Bacalini et al. (2015) [37] and Ghezzo et al. (2014) [38] provided further details regarding ethical approval, data collection, DNA extraction and DNAm measurement (including probe exclusion). The Werner’s syndrome dataset (GSE131752) was taken from the following studies [16, 24] and all details regarding the study population and data set can be found there. The DNAm dataset (cases *n*=24 and matched controls *n*=24) was for the EPIC array consisting of 850K probes as opposed to the 450K probes used in the DNAm datasets for the other analyses.

The DNAm MoBa2 (*n*=685) data was measured using the Illumina HumanMethylation450 BeadChip (Illumina, San Diego, CA, USA). The quality control process has been described previously [25]. The prenatal smoking variable was defined as described in Joubert and colleagues’ previous publication [27] but only for “sustained smoking” (*n*=70) and “never smoked” (*n*=512). Maternal BMI (15.2, 43.2) was a continuous trait available for *n*=657 out of 685 samples.

## LIST ABBREVIATIONS

BMI: Body mass index
CA: Chronological age
CpG: 5’-C-phosphate-G-3’,CG dinucleotide
DNAm: DNA methylation
EA: Epigenetic age
EAA: Epigenetic age acceleration
eEAAn: Epigenetic age acceleration related principal component *n*
eEGAAn: Epigenetic gestational age acceleration related principal component *n*
EGAA: Epigenetic gestational age acceleration
EWAS: Epigenome-Wide Association Study
GA: Gestational age
MoBa: The Norwegian Mother, Father and Child Cohort Study
P_B_: Bonferroni adjusted p-value
PCA: Principal component analysis
PC: Principal component

## DECLARATIONS

### Ethics approval and consent to participate

The MoBa study was approved by the Regional Committee for Ethics in Medical Research, the Norwegian Data Inspectorate, and the Institutional Review Board of the National Institute of Environmental Health Sciences, USA. The experimental methods comply with the Helsinki Declaration.

### Consent for publication

Written informed consent was provided by all mothers participating in the MoBa-study.

### Availability of data and materials

Two of the datasets (GSE52588, GSE131752) supporting the conclusions of this article is publically available in Gene Expression Omnibus repository, https://www.ncbi.nlm.nih.gov/geo/. MoBa2 can be accessible upon an application to Norwegian Institute of Public Health (NIPH, http://www.fhi.no/en/).

### Authors’ contributions

YL and JB contributed equally to the method, writing, computation and initiation of the project.

## Acknowledgements

We appreciate all the study participants in GSE52588 and GSE131752 as well as the Norwegian Mother and Child Cohort Study. We also thank all the researchers who made the datasets GSE52588 and GSE131752 publically available.

## ADDITIONAL FILES

**Additional file 1**

Contains an R script file with step-by-step instructions on how to perform the proposed procedure

**Additional file 2**

Contains a .docx file figures S1 and S2 with explanations of the Werner’s syndrome data.

**Additional file 3**

Contains a .docx file of loadings for eEGAA9 and eEGAA90 (maternal smoking). CpGs found to also be associated with maternal smoking in a meta-analysis [27] are highlighted.

**Additional file 4**

Contains a .docx file Figure S4, a visualization of the eE(G)AA extraction procedure

## REFERENCES

1. Koch CM, Wagner W: Epigenetic-aging-signature to determine age in different tissues. Aging (Albany NY) 2011, 3(10):1018–1027.

2. Hannum G, Guinney J, Zhao L, Zhang L, Hughes G, Sadda S, Klotzle B, Bibikova M, Fan JB, Gao Y et al: Genome-wide methylation profiles reveal quantitative views of human aging rates. Mol Cell 2013, 49(2):359–367.

3. Horvath S: DNA methylation age of human tissues and cell types. Genome biology 2013, 14(10):R115.

4. Bocklandt S, Lin W, Sehl ME, Sanchez FJ, Sinsheimer JS, Horvath S, Vilain E: Epigenetic predictor of age. PloS one 2011, 6(6):e14821.

5. Marioni RE, Shah S, McRae AF, Chen BH, Colicino E, Harris SE, Gibson J, Henders AK, Redmond P, Cox SR et al: DNA methylation age of blood predicts all-cause mortality in later life. Genome biology 2015, 16:25.

6. Levine AJ, Quach A, Moore DJ, Achim CL, Soontornniyomkij V, Masliah E, Singer EJ, Gelman B, Nemanim N, Horvath S: Accelerated epigenetic aging in brain is associated with pre-mortem HIV-associated neurocognitive disorders. J Neurovirol 2016, 22(3):366–375.

7. Levine ME, Lu AT, Chen BH, Hernandez DG, Singleton AB, Ferrucci L, Bandinelli S, Salfati E, Manson JE, Quach A et al: Menopause accelerates biological aging. Proc Natl Acad Sci U S A 2016, 113(33):9327–9332.

8. Horvath S, Ritz BR: Increased epigenetic age and granulocyte counts in the blood of Parkinson’s disease patients. Aging (Albany NY) 2015, 7(12):1130–1142.

9. Horvath S, Oshima J, Martin GM, Lu AT, Quach A, Cohen H, Felton S, Matsuyama M, Lowe D, Kabacik S et al: Epigenetic clock for skin and blood cells applied to Hutchinson Gilford Progeria Syndrome and ex vivo studies. Aging (Albany NY) 2018, 10(7):1758–1775.

10. Levine ME, Lu AT, Bennett DA, Horvath S: Epigenetic age of the pre-frontal cortex is associated with neuritic plaques, amyloid load, and Alzheimer’s disease related cognitive functioning. Aging (Albany NY) 2015, 7(12):1198–1211.

11. Horvath S, Langfelder P, Kwak S, Aaronson J, Rosinski J, Vogt TF, Eszes M, Faull RL, Curtis MA, Waldvogel HJ et al: Huntington’s disease accelerates epigenetic aging of human brain and disrupts DNA methylation levels. Aging (Albany NY) 2016, 8(7):1485–1512.

12. Raina A, Zhao X, Grove ML, Bressler J, Gottesman RF, Guan W, Pankow JS, Boerwinkle E, Mosley TH, Fornage M: Cerebral white matter hyperintensities on MRI and acceleration of epigenetic aging: the atherosclerosis risk in communities study. Clin Epigenetics 2017, 9:21.

13. Horvath S, Raj K: DNA methylation-based biomarkers and the epigenetic clock theory of ageing. Nature reviews Genetics 2018, 19(6):371–384.

14. Field AE, Robertson NA, Wang T, Havas A, Ideker T, Adams PD: DNA Methylation Clocks in Aging: Categories, Causes, and Consequences. Mol Cell 2018, 71(6):882–895.

15. Horvath S, Garagnani P, Bacalini MG, Pirazzini C, Salvioli S, Gentilini D, Di Blasio AM, Giuliani C, Tung S, Vinters HV et al: Accelerated epigenetic aging in Down syndrome. Aging Cell 2015, 14(3):491–495.

16. Maierhofer A, Flunkert J, Oshima J, Martin GM, Haaf T, Horvath S: Accelerated epigenetic aging in Werner syndrome. Aging (Albany NY) 2017, 9(4):1143–1152.

17. Horvath S, Pirazzini C, Bacalini MG, Gentilini D, Di Blasio AM, Delledonne M, Mari D, Arosio B, Monti D, Passarino G et al: Decreased epigenetic age of PBMCs from Italian semi-supercentenarians and their offspring. Aging (Albany NY) 2015, 7(12):1159–1170.

18. Girchenko P, Lahti J, Czamara D, Knight AK, Jones MJ, Suarez A, Hamalainen E, Kajantie E, Laivuori H, Villa PM et al: Associations between maternal risk factors of adverse pregnancy and birth outcomes and the offspring epigenetic clock of gestational age at birth. Clin Epigenetics 2017, 9:49.

19. Knight AK, Craig JM, Theda C, Baekvad-Hansen M, Bybjerg-Grauholm J, Hansen CS, Hollegaard MV, Hougaard DM, Mortensen PB, Weinsheimer SM et al: An epigenetic clock for gestational age at birth based on blood methylation data. Genome biology 2016, 17(1):206.

20. Zhang Q, Vallerga CL, Walker RM, Lin T, Henders AK, Montgomery GW, He J, Fan D, Fowdar J, Kennedy M et al: Improved precision of epigenetic clock estimates across tissues and its implication for biological ageing. Genome Med 2019, 11(1):54.

21. Khouja JN, Simpkin AJ, O’Keeffe LM, Wade KH, Houtepen LC, Relton CL, Suderman M, Howe LD: Epigenetic gestational age acceleration: a prospective cohort study investigating associations with familial, sociodemographic and birth characteristics. Clin Epigenetics 2018, 10:86.

22. Tibshirani R: Regression shrinkage and selection via the lasso. Journal of the Royal Statistical Society: Series B (Methodological) 1996, 58(1):267–288.

23. Quach A, Levine ME, Tanaka T, Lu AT, Chen BH, Ferrucci L, Ritz B, Bandinelli S, Neuhouser ML, Beasley JM et al: Epigenetic clock analysis of diet, exercise, education, and lifestyle factors. Aging (Albany NY) 2017, 9(2):419–446.

24. Maierhofer A, Flunkert J, Oshima J, Martin GM, Poot M, Nanda I, Dittrich M, Muller T, Haaf T: Epigenetic signatures of Werner syndrome occur early in life and are distinct from normal epigenetic aging processes. Aging Cell 2019, 18(5):e12995.

25. Bohlin J, Haberg SE, Magnus P, Reese SE, Gjessing HK, Magnus MC, Parr CL, Page CM, London SJ, Nystad W: Prediction of gestational age based on genome-wide differentially methylated regions. Genome biology 2016, 17(1):207.

26. Simpkin AJ, Suderman M, Howe LD: Epigenetic clocks for gestational age: statistical and study design considerations. Clin Epigenetics 2017, 9:100.

27. Joubert BR, Felix JF, Yousefi P, Bakulski KM, Just AC, Breton C, Reese SE, Markunas CA, Richmond RC, Xu CJ et al: DNA Methylation in Newborns and Maternal Smoking in Pregnancy: Genome-wide Consortium Meta-analysis. American journal of human genetics 2016.

28. Teschendorff AE, West J, Beck S: Age-associated epigenetic drift: implications, and a case of epigenetic thrift? Human molecular genetics 2013, 22(R1):R7–R15.

29. Sandoval J, Heyn H, Moran S, Serra-Musach J, Pujana MA, Bibikova M, Esteller M: Validation of a DNA methylation microarray for 450,000 CpG sites in the human genome. Epigenetics: official journal of the DNA Methylation Society 2011, 6(6):692–702.

30. Wu MC, Joubert BR, Kuan PF, Haberg SE, Nystad W, Peddada SD, London SJ: A systematic assessment of normalization approaches for the Infinium 450K methylation platform. Epigenetics: official journal of the DNA Methylation Society 2014, 9(2):318–329.

31. Engebretsen S, Bohlin J: Statistical predictions with glmnet. Clin Epigenetics 2019, 11(1):123.

32. Martin-Herranz DE, Aref-Eshghi E, Bonder MJ, Stubbs TM, Choufani S, Weksberg R, Stegle O, Sadikovic B, Reik W, Thornton JM: Screening for genes that accelerate the epigenetic aging clock in humans reveals a role for the H3K36 methyltransferase NSD1. Genome biology 2019, 20(1):146.

33. Chen BH, Marioni RE, Colicino E, Peters MJ, Ward-Caviness CK, Tsai PC, Roetker NS, Just AC, Demerath EW, Guan W et al: DNA methylation-based measures of biological age: meta-analysis predicting time to death. Aging (Albany NY) 2016, 8(9):1844–1865.

34. Yohai V, Stahel W, Zamar R: A Procedure for Robust Estimation and Inference in Linear Regression. In., vol. 34: Springer New York; 1991: 365–374.

35. Friedman J, Hastie T, Tibshirani R: Regularization paths for generalized linear models via coordinate descent. Journal of statistical software 2010, 33(1):1.

36. Team RC: R: A language and environment for statistical computing. R foundation for Statistical Computing 2005.

37. Bacalini MG, Gentilini D, Boattini A, Giampieri E, Pirazzini C, Giuliani C, Fontanesi E, Scurti M, Remondini D, Capri M et al: Identification of a DNA methylation signature in blood cells from persons with Down Syndrome. Aging (Albany NY) 2015, 7(2):82–96.

38. Ghezzo A, Salvioli S, Solimando MC, Palmieri A, Chiostergi C, Scurti M, Lomartire L, Bedetti F, Cocchi G, Follo D et al: Age-related changes of adaptive and neuropsychological features in persons with Down Syndrome. PLoS One 2014, 9(11):e113111.

