## Supplementary information 1 for "Amplification of the epigenetic (gestational) age acceleration signal"

```
# Need packages glmnet and parallel:
require(parallel)
require(glmnet)

# Script for eE(G)AA procedure
# This is for demonstration only as it must be modified to fit different
datasets

# Simple imputation for missing gestational age values (GA_MoBa_2)
temp <- GA_MoBa_2
temp[which(is.na(temp))]<-median(temp, na.rm=T)
GA_MoBa_2_IMP <- temp

# Make residual matrix for Bohlin clock
# Clock CpGs ncol=96, samples nrow=685
resMatr<-matrix(NA, ncol=96, nrow=685)
rownames(resMatr) <- 1:685
# Rename columns to Bohlin clock CpG's found in the CpG matrix GA_CpGs
colnames(resMatr) <- colnames(GA_CpGs)
# Perform linear regression for gestational age on each of the 96 clock
CpG's
rNames<-NA
for(i in 1:96){
    mod<-lm( GA_MoBa_2_IMP ~ GA_CpGs[,i] )
    rNames<-as.integer(names(resid(mod)))
    resMatr[rNames,i] <- as.vector(resid(mod))
}
# The matrix should now contain 96 columns corresponding to the error
residuals in each of the 96 regressions

# Perform PCA on the residual matrix
resMatrPCA <- prcomp( resMatr )

# Checking out maternal smoking on MoBa, can not include missing
# pDat$smk2yn is the MoBa 2 smoking variable (binary yes/no)
temp<-pDat$smk2yn
temp2<-temp[which(!is.na(temp))]
pcaMatrTemp<-resMatrPCA$x[which(!is.na(temp)),]

# Lasso on maternal smoking (outcome) with respect to PCA matrix
# Might have to do this multiple times if nothing is found (due to the
cross-validation procedure)
# It's recommended that a seed is being set (set.seed())

# Binary outcome must be a factor
temp2<-as.factor(temp2)

# Perform cross validates Lasso in parallel
mod.cv <- cv.glmnet( y = temp2, x = pcaMatrTemp, family="binomial",
parallel = T )

# Pick out components based on minimal penalty
tempMat=as.matrix(coef(mod.cv, s="lambda.min"))
# Arrange components in a data frame
tempMat=as.data.frame(tempMat)
names(tempMat) <- "beta"
tempMat$PC <- rownames(tempMat)
# Include only non-zero eEGAA's with non-zero slope-parameter
```

```
res_pc = tempMat[which(tempMat$beta!=0.00),]
# Remove intercept
res_pc <- res_pc[-1,]
# A binomial regression can now be carried out based on extracted eEGAA's
mod<-glm( temp2 ~ pcaMatrTemp[,res_pc$PC], family=binomial)
AIC(mod)
# And compared too a null-model
mod0<-glm( temp2 ~ 1, family=binomial)
AIC(mod0)

# Loadings can also be examined to interrogate CpG's for 'extractedCpGs'
resMatrPCA$rotation[,extractedCpGs]
```
