## Supplementary information 2 for "Amplification of the epigenetic (gestational) age acceleration signal"

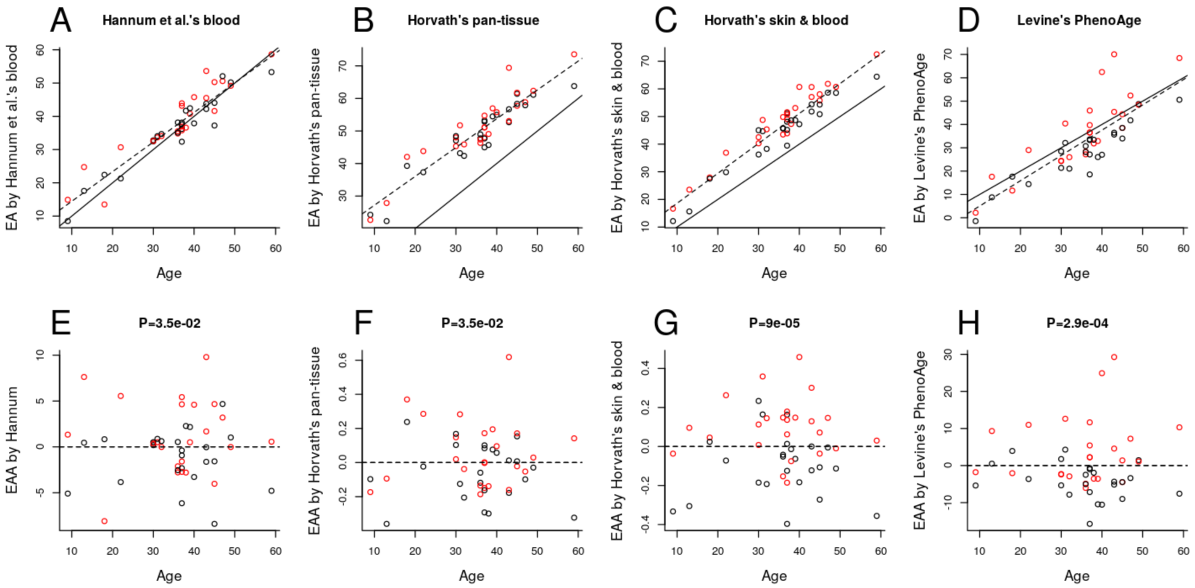


**Figure S1. Associations between residual EAA and Werner's syndrome.**

**A-D**: Scatter plot of CA versus EA. **E-H**: Scatter plot of CA versus EAA.

The black dots refer to unaffected controls whereas the red dots refer to subjects with Werner's syndrome. P-values were obtained from student t tests comparing EAAs between controls and subjects with Werner's syndrome.

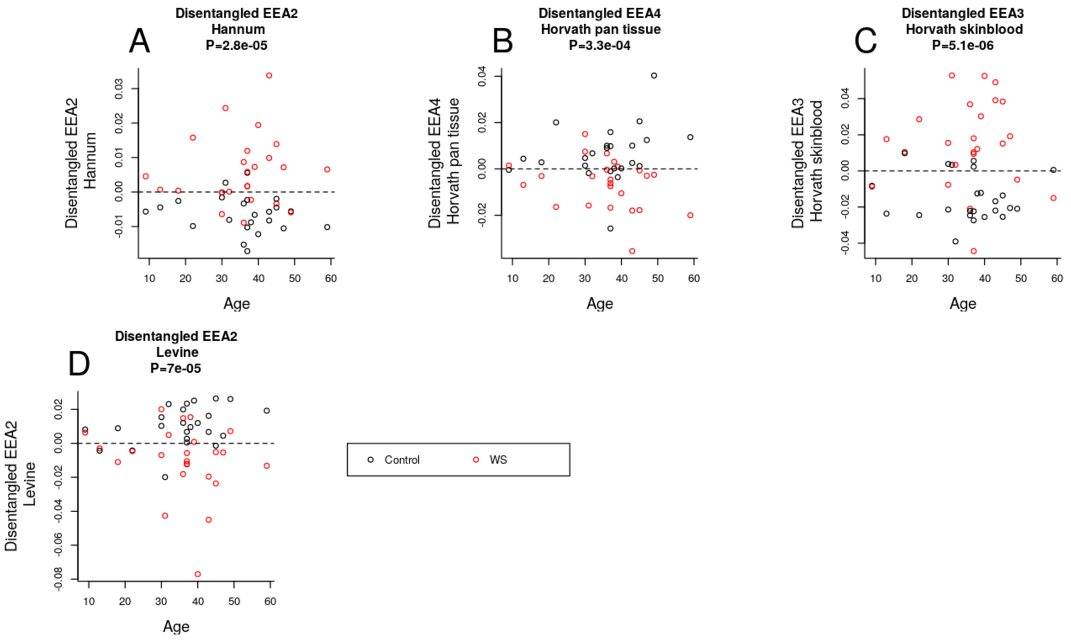


**Figure S2. Associations between eEAAs and Werner's syndrome.**

**A:** eEAA2 (i.e., PC2) obtained by Hannum et al.'s clock. **B:** eEAA4 (i.e., PC4) obtained by Horvath's pan-tissue clock. **C:** eEAA3 (i.e., PC3) obtained by Horvath et al.'s skin&blood clock. **D:** eEAA2 (i.e., PC2) obtained by Levine et al.'s PhenoAge clock.

The black dots refer to unaffected controls whereas the red dots refer to subjects with Werner's syndrome. P-values were obtained from student t tests comparing eEAAs between controls and subjects with Werner's syndrome.
