## Supplementary information 3 for "Amplification of the epigenetic (gestational) age acceleration signal"

| | eEGAA9 | eEGAA90 |
|---|---|---|
| cg00153101 | 0.117747540 | -1.202500e-02 |
| ***cg00602416*** | ***0.019454336*** | ***1.793465e-02*** |
| cg00711496 | 0.029077775 | 1.313746e-02 |
| cg01190109 | 0.023553166 | -1.035688e-01 |
| cg01281797 | 0.019938244 | -9.894078e-04 |
| cg01635555 | 0.042272843 | -7.924856e-03 |
| ***cg01833485*** | ***-0.022142815*** | ***-2.186181e-03*** |
| cg02324006 | 0.025583726 | -1.202360e-02 |
| cg02405476 | -0.080291963 | 7.010789e-03 |
| cg02567958 | 0.017656597 | 3.126835e-05 |
| cg02642822 | 0.020794799 | -1.329667e-03 |
| cg03098721 | -0.056497976 | -1.750730e-03 |
| cg03108070 | -0.293644547 | -5.609284e-03 |
| cg03281561 | -0.115243120 | 1.265927e-02 |
| cg03337084 | -0.050117483 | 2.222490e-03 |
| cg03507326 | -0.026704228 | 6.024120e-03 |
| cg03710860 | 0.031169328 | -4.060238e-02 |
| cg03729251 | -0.041832548 | 3.161857e-04 |
| ***cg03773820*** | ***0.072592037*** | ***-2.792355e-03*** |
| cg03963689 | 0.059428680 | -2.359357e-03 |
| cg04347477 | -0.014584367 | 6.050106e-03 |
| cg04685228 | -0.026111510 | -7.309373e-03 |
| cg05053327 | 0.244461212 | -2.194024e-03 |
| cg05544807 | -0.005117413 | 3.449874e-02 |
| cg05877497 | -0.235680704 | -8.271426e-03 |
| ***cg06753281*** | ***0.030207814*** | ***1.155253e-02*** |
| cg06897661 | 0.127381217 | -2.692847e-03 |
| cg07106169 | -0.099440900 | -8.656554e-03 |
| cg07676709 | 0.015295054 | -3.841453e-02 |
| ***cg07738730*** | ***0.259690972*** | ***4.412407e-03*** |
| cg07749613 | -0.040853906 | -3.701967e-03 |
| cg07788865 | 0.016710908 | -1.115363e-02 |
| cg07816074 | -0.048052498 | -4.611667e-03 |
| cg07835443 | 0.001496119 | -3.688585e-03 |
| cg08326019 | -0.008783222 | -1.495589e-02 |
| cg08620426 | 0.040719803 | -3.138119e-03 |
| cg08943494 | -0.022149291 | 4.887351e-03 |
| cg08965235 | 0.048741004 | 4.024634e-04 |
| cg09447786 | -0.090602841 | -5.634393e-03 |
| cg10107671 | 0.019924664 | -4.204114e-02 |
| cg10308785 | 0.004454848 | 4.140828e-03 |
| cg11124260 | -0.006213392 | 3.126568e-03 |
| cg11294761 | -0.050435086 | 3.001295e-03 |
| ***cg11864574*** | ***0.022639602*** | ***-2.209746e-02*** |
| cg12564453 | 0.029232136 | 2.084225e-02 |
| cg12880227 | 0.022064079 | 6.222212e-03 |
| cg12999267 | -0.108596638 | 2.324043e-03 |
| cg13036381 | -0.082607056 | -2.658200e-03 |
| cg13066703 | 0.016290622 | -6.037007e-04 |

| | | |
|---|---|---|
| cg13433246 | -0.063380373 | 6.431916e-03 |
| cg13641317 | -0.098467915 | 6.108256e-03 |
| cg13733403 | -0.007353530 | 1.113873e-02 |
| cg13959344 | 0.001106577 | 2.538499e-02 |
| cg13982823 | 0.019540653 | -3.507479e-03 |
| cg14276580 | -0.187796660 | 1.733768e-04 |
| cg14427590 | -0.005764658 | 2.819953e-02 |
| cg15035133 | 0.027955737 | -1.929062e-03 |
| cg15131146 | -0.069636260 | -1.530457e-03 |
| cg15165154 | 0.012988224 | -1.547819e-03 |
| ***cg15626350*** | ***0.010742062*** | ***1.900190e-02*** |
| ***cg15908709*** | ***0.042764602*** | ***7.169300e-02*** |
| cg16187883 | 0.018288461 | 7.663461e-01 |
| ***cg16348385*** | ***-0.109631271*** | ***-5.868935e-04*** |
| cg16536918 | 0.035083670 | 2.354769e-02 |
| cg17022232 | 0.033320570 | -8.471454e-03 |
| ***cg18183624*** | ***0.441710361*** | ***1.656837e-03*** |
| cg18217136 | -0.218524857 | 5.464515e-04 |
| cg18954401 | -0.063575872 | -1.551583e-02 |
| ***cg19057830*** | ***-0.018585113*** | ***3.989232e-03*** |
| cg19439123 | 0.028306991 | -7.342263e-03 |
| cg19875532 | 0.073008399 | -3.619352e-03 |
| cg20301308 | -0.020085159 | 8.085235e-03 |
| cg20303561 | -0.003591649 | -2.899820e-02 |
| cg20377955 | 0.050003355 | -1.955752e-02 |
| ***cg20816447*** | ***0.039416962*** | ***8.819980e-03*** |
| cg21081878 | -0.274326547 | -2.715845e-03 |
| cg21143441 | 0.021395806 | 6.467264e-03 |
| cg21155834 | 0.030515356 | 3.196434e-04 |
| cg21221899 | 0.329090612 | -1.218958e-02 |
| cg21707172 | 0.005484062 | -2.332052e-02 |
| cg21878650 | 0.030204665 | -1.007372e-03 |
| cg22761205 | -0.025331052 | 2.164137e-03 |
| cg22796593 | 0.019669809 | -1.153278e-03 |
| cg22797644 | 0.046005641 | 4.002320e-02 |
| cg23051248 | -0.056031216 | -6.657283e-03 |
| cg23346945 | 0.023712870 | 5.081834e-02 |
| cg23403099 | 0.025095083 | -3.655530e-02 |
| cg23457357 | 0.002615428 | -1.594606e-02 |
| ***cg24041556*** | ***0.112868498*** | ***-4.924879e-03*** |
| ***cg24087613*** | ***0.017730803*** | ***-6.126367e-01*** |
| cg24366564 | -0.243053113 | -1.584424e-03 |
| ***cg25150953*** | ***0.087095830*** | ***-2.809881e-03*** |
| cg25531857 | 0.038423553 | -3.561450e-02 |
| cg25639749 | -0.017058853 | 2.407693e-03 |
| cg26077811 | 0.007016001 | -9.887147e-03 |
| cg26092675 | -0.003810519 | -9.983727e-03 |
