## Supplementary information 4 for "Amplification of the epigenetic (gestational) age acceleration signal"

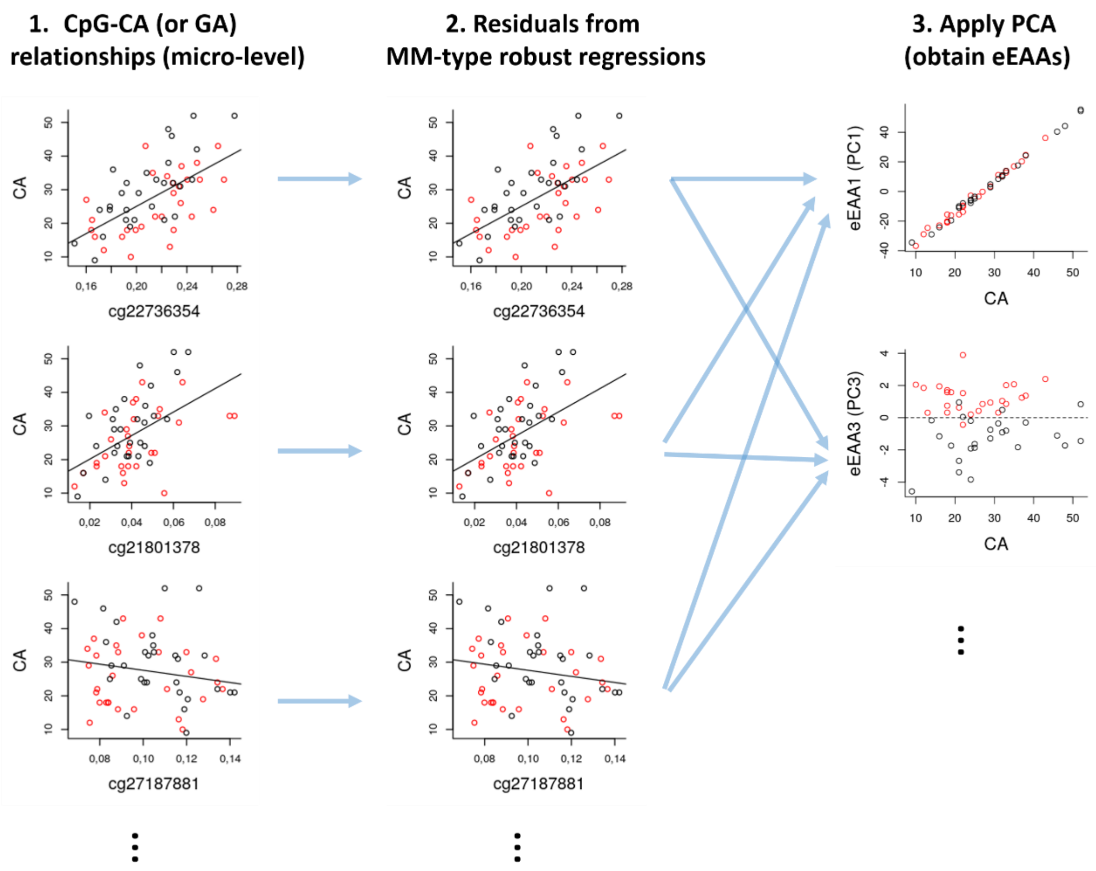


**Figure S4. Visualization of the E(G)AA-extraction method.**

The CpGs were selected by Levine’s PhenoAge clock. For better understanding, we visualized CpG-CA relationships where subjects with Down syndrome (red dots) formed relatively clear shifts.
